# Decision tree models and cell fate choice

**DOI:** 10.1101/2020.12.19.423629

**Authors:** Ivan Croydon Veleslavov, Michael P.H. Stumpf

## Abstract

Single cell transcriptomics has laid bare the heterogeneity of apparently identical cells at the level of gene expression. For many cell-types we now know that there is variability in the abundance of many transcripts, and that average transcript abundance or average gene expression can be a unhelpful concept. A range of clustering and other classification methods have been proposed which use the signal in single cell data to classify, that is assign cell types, to cells based on their transcriptomic states. In many cases, however, we would like to have not just a classifier, but also a set of interpretable rules by which this classification occurs. Here we develop and demonstrate the interpretive power of one such approach, which sets out to establish a biologically interpretable classification scheme. In particular we are interested in capturing the chain of regulatory events that drive cell-fate decision making across a lineage tree or lineage sequence. We find that suitably defined decision trees can help to resolve gene regulatory programs involved in shaping lineage trees. Our approach combines predictive power with interpretabilty and can extract logical rules from single cell data.

## 1 Introduction

Trees have been providing an organising framework for biology since before Darwin and Wallace. Descent from a single common ancestor also provides an organising principle in developmental biology(*1*–*4*). The development of multicellular organisms appears to be an astonishingly well choreographed process. This involves a sequence of transitions starting from omnipotent cells, via multi-potent and progenitor cells, all the way to terminally differentiated cells. From a single fertilised egg cell several trillion cells derive in humans. In *Caenorhabditis celegans* female the developmental tree depicting the relationships among all 959 somatic cells in the worm adult have been mapped painstakingly by John Sulston and colleagues (*5*).

Technological advances are now allowing us to watch the developmental process unfold in great detail in other organisms. Experiments in *Danio rerio* (*6*) and *Xenopus* (*7*) have traced cell lineage trees from the fertilised egg cell for several generations. Further analyses in gastruloids and organoids provide clues about similar dynamics in experimentally convenient models for other organisms (*8*–*10*). From these observations, and now the ability to assay gene expression pattern in single cells(*11*), we can start to learn regulatory programs underlying developmental processes.

Here we develop a decision-tree framework to learn developmental programs of cell-fate choice from single cell data. Decision trees represent sequences of binary choices; going through this sequence assigns data to labels. In the current context labels can represent cell-fates or phenotypically distinct cell types, and the choices to be made relate to changes in expression levels of certain genes (*11, 12*). Which genes, and which levels are discriminatory is determined by the algorithm described below in detail. There are two main advantages of the resulting tree: (i) it is directly interpretable, and from the tree we can distill mechanistic hypotheses that are testable as well as predictive; (ii) learning such trees is feasible even for very large data-sets with low computational effort, one of the fundamental challenges in this field (*12*).

Classification of cells based on their transcriptomic signatures can be performed in a number of different ways (see for example (*13*–*15*)). Deep learning methods have become particularly popular; their main advantages are that they are capable of learning feature sets used for classification and (locally) optimal classifiers with minimal intervention (*16*). They do, however, require vast amounts of data, and despite their computational sophistication, are computationally costly. Other, more conventional classification methods (*17, 18*) – logistic regression, support vector machines, kernel-based classifiers, … – can also be applied. And the predictive power of decision tress can be improved through random forests.

Classification typically involves a trade-off between the predictive performance and the interpretability of classifiers. The choice of which classifier to use depends, among many other factors, on the data, available background knowledge, but also the objectives of the investigators. No single method can be optimal under every condition (*19*). Decision trees are on the “interpretable end of the spectrum” of classification methods, and here we develop this in order to develop better insights into the transcriptional events underlying e.g. cell differentiation (*3, 20*).

Naturally, we cannot expect the decision tree to mirror the tree (or, to be precise, the state-transition diagram) faithfully capturing the relationships between cell-types (*1, 2*). Cell-fates differ by more than the expression in a small number of genes, whereas most decision trees use simple binary choices at the branching points. Below we will show that there is a high-level of concordance between suitably defined decision trees and cell trees. But rather than merely recapitulating the developmental process, the decision tree constitutes a simple and mechanistically interpretable model of this process.

Our approach sets out to construct trees such that each split in the tree yields maximal information about the population sample of cells. We construct trees based on this criterion, and show that they are capable of capturing known biological relationships and we rediscover sets of genes known to be pivotal for different cell differentiation processes. But we also use the tree as a tool to understand potential ways of guiding these processes.

## 2 Results

### 2.1 Learning decision tree models from single-cell data

#### 2.1.1 Mouse neuronal precursors

To demonstrate how decision tree models can be used to characterise developmental boundaries from single-cell transcriptomic data (*20, 21*), we train a decision tree model using an implementation of the Classification and Regression Tree (CART) algorithm (see 4.1.1) on a murine data set for the cell stage target variable on a random 80% training partition. The resulting model grows 22 internal nodes, and 23 leaf nodes to distinguish the ESC, EPI, and NPC labels in the training data, Figure 3a. Applying the learnt model to the withheld test data, this decision tree correctly classifies 113 of the 129 withheld cells (87.5% accuracy).

We also perform 5-fold cross validation on the `Murine` data for the cell stage target variable. restricting the trees to a maximum depth of four nodes yields an average predictive performance of 95.6% across the folds. The sum confusion matrix for this cross validation is shown, see Table 1. Reassuringly, we observe that no sample with true label ‘ES’ is labelled as ‘NPC’ and *vice versa*. Indeed, all misclassifications occur at the EPI/NPC boundary – with 1.16% of NP cells being misclassified as EPI and 1.41% of EPI cells being misclassified as NP cells. This demonstrates the ability of decision trees to capture latent biological signals from single-cell transcriptomic data: the model learnt during training – without *a priori* constraints – clearly and correctly captures that cells progress from ESC to EPI to NPC stages.

**Table 1:** Confusion matrix for decision trees learnt on the `Murine` data for the cell stage target variable, summed across all folds of 5-fold cross-validation

|  |  | Predicted |  |  | Total |
| --- | --- | --- | --- | --- | --- |
|  |  | ESC | EPI | NPC |  |
| Actual | ESC | 94 | 0 | 0 | 94 |
|  | EPI | 0 | 210 | 3 | 213 |
|  | NPC | 0 | 3 | 235 | 238 |
| Total | | 94 | 213 | 238 | $N = 545$ |

In addition, we provide Sankey diagram representation for a tree grown on a random 80% of the data to highlight the flow of training samples through the internal nodes, Figure 3b. In this representation the width of the bars is proportional to the weight (or number) of samples that take that route through the tree, and the tone of the internal branches corresponds to the entropy over the constituent cell labels. We see that much of the model’s branches are involved in separating ES and EPI cells from each other, as well as EPI cells and NPC. The majority of the separation of ES and NP cells is achieved by the first branching point, which splits the samples based on whether *Fgf4* expression is greater than 1.9 (dimensionless relation measured during qPCR)(*20*).

### 2.2 Knowledge discovery

#### 2.2.1 Classifying unlabelled samples

Given a cell with unknown class label, a learnt decision tree model allows for fast and interpretable sample classification. The unlabelled sample is simply passed through the tree, starting at the topmost node. The sample then takes a path through the internal nodes according to the constituent features and thresholds until the unlabelled sample reaches a terminal (leaf) node. Here, the unlabelled sample is classified using the mode label of samples present in that terminal node from model training.

The classification of unlabelled samples is thus fast as it corresponds to a series of binary splits represented in Boolean conditional logic (*22, 23*). Importantly, the ‘reasoning’ behind the label attributed by the model is also immediately interpretable if one keeps track of not simply the label assigned by the model, but also of the path taken by the sample through the tree.

Further, we might consider that sub-samples of cell classes that share a specific terminal leaf may in general be more similar than those that were attributed the same class, but by a different terminal node and thus a different path through the learnt model. This could be used to investigate heterogeneity within cell stages, or to probe the existence of (cryptic) sub-populations within established cell classes.

#### 2.2.2 Extracting Boolean rule sets

With a decision tree in hand it also becomes straightforward to extract Boolean rules governing classification boundaries (*22*). Iterating through all paths from the root node in the model through to the leaf nodes provides a set of rules learnt by the model during training to allocate samples to classes. If we are confident in our model’s ability to describe the system well – for example, upon seeing good performance in generalisation to unseen data – the learnt rules can therefore be used as a base for knowledge discovery and mechanistic hypothesis generation (*23, 24*). We propose that, for instance, the minimal rules learnt in the context of cell development could be used directly in the formulation of a simple, low-dimensional, mechanistic model of the cell stage transition. Whilst we would not expect such a model to capture the full complexity of the underlying cellular and transcriptomic processes, we posit that this decision-tree-led approach offers considerable value, especially in contexts where high-dimensional data and limited knowledge *a priori* would otherwise prevent easy intuition. Additionally, these Boolean rules could be used to design a combinatorial system of differentiation markers for experimental use e.g. for engineering cohorts of FACS cell surface markers.

The explicit rule set learnt by the model discussed in section 2.1.1 is given in concrete Boolean terms in the supplementary material (section A.1).

#### 2.2.3 Designing minimal perturbations for sample reclassification

One major motivation for developing tree-based models of cell fate commitment is to aid the design of cell perturbations that can guide cell fate decisions (*25*–*27*). As already demonstrated, decision trees learn a set of features and associated thresholds to classify samples. The path a given sample takes through the tree is thus defined by the sample’s expression levels for those learnt features relative to the corresponding learnt thresholds. Given a learnt decision tree model and a sample it is clear to see how changing the expression of the sample for those embedded features may alter the path taken by the sample and thus its classification by the model. Explicitly, we can exploit the structure of the learnt classifier in order to design minimal perturbations that when applied to a sample will result in its reclassification to a target label of our choosing.

To do so, we have developed a flexible algorithm that takes a learnt decision model, a prospective sample and a target cell class, and returns the list of minimal changes that, if applied to the sample would result in the system moving to the desired target class (see methods, section 4.3). Here we apply this scheme to the same decision tree model learnt in section 2.1.1. Sample #10 from the test set is classified as an ESC. If we want to drive a cell from the same starting point towards, for example, the EPI target fate, we are able to generate all possible, minimal sets of potential changes that will cause the decision tree to assign sample #10 to the EPI class [Table 2]. Ranking these candidate sets of proposed changes by total absolute tree path distance (*28*), we can see that the smallest change that results in the sample reclassification is to up-regulate *Mef2a* and down-regulate *Klf4*. Note there are 12 sets of changes generated by the algorithm, corresponding to the 12 EPI terminal nodes in the learnt model; and thus there are 12 unique paths that can result in the sample being classified as EPI. Applying any of these perturbations to the sample and feeding the adjusted sample to the model results in the expected reclassification to the EPI class. The framework here can thus support *in vivo* synthetic biology and regenerative medicine, by rationally proposing interventions and control measures that affect cell fate.

**Table 2:**
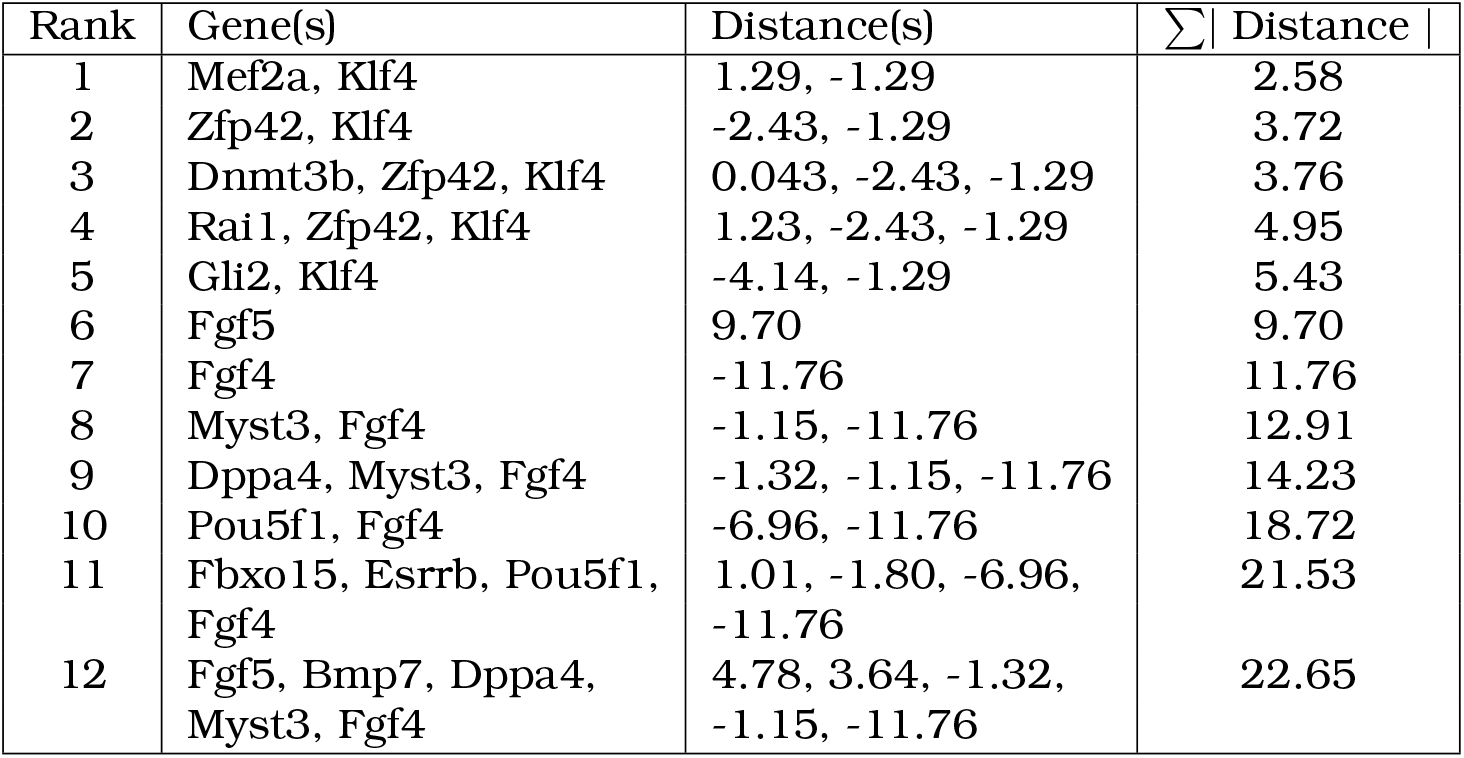
Minimum perturbations that result in the reclassification of test sample #10 by the learnt model from its initial classification as ESC to its target classification of EPI. The 12 unique paths to reclassification are ranked by the sum of absolute distances.

### 2.3 Incorporating known lineage hierarchies

Existing knowledge of lineage hierarchies can be incorporated into our decision tree framework. This can aid the discovery of new rules from single cell transcriptomic data that are commensurate with, e.g. available lineage information.

Using a previously established developmental lineage hierarchy for *Xenopus tropicalis* as a scaffold, we grow decision trees at each known branching point in the lineage map. For each branching point, from stage-08 through to stage-14, we filter the data for samples with the branch-relevant cell stage target labels. The individual trees learnt on these filtered subsets, all of which are limited to a depth of six nodes, are then assembled over the known lineage hierarchy. The result is a detailed lineage map of development in the early *Xenopus tropicalis* embryo at the transcriptomic level, with the most informative genes for each decision boundary learnt directly from the data *a priori* with decision tree classifiers, Figure 4.

From the resulting lineage-map/decision-tree hybrid, we can trace a cell from the earliest cell type, ‘Stage 8 Blastula’, through the learnt portions of the tree to the most mature ‘Stage 14’ cell types; ‘chordal neural plate border’,’migrating myeloid progenitors’, ‘presomitic mesoderm’ *etc*. Critically, at each branching point we can directly observe the features learnt during the training process that are the most informative for characterising each of those transitions.

This new approach of using decision tree classifiers in conjunction with known lineage maps offers the possibility to meld developmental biology knowledge with single-cell transcriptomic information. As large scale ATLAS projects (*29*) continue to generate vast data sets across developmental processes and unique tissues, we see this approach as holding significant promise for delineating the complex genetic interactions underpinning e.g. ontogenesis and self-renewal(*3*).

## 3 Discussion

There is pressing need in genome and developmental biology for mechanistic models that make effective use of single-cell transcriptomic data to characterise complex developmental processes (*12*). Our hope is that through careful design choices, the use of these models may extend beyond simply classifying unlabelled samples with high performance and rather that we might exchange small trade-offs in predictive performance for an improved understanding of the underlying system.

Here, we showed how suitably constructed decision tree models go some way to bridging this gap. Training decision tree models on single-cell transriptomic data from RT-qPCR and sc-RNAseq allows us to characterise murine neuronal differentiation as well as early *Xenopus tropicalis* embryogenesis. We demonstrate that these classifiers perform well on withheld data and show how the structure of the learnt models also allows for feature selection, Boolean rule-set generation and for the design of perturbations for targeted sample reclassification.

The results we showcase here represent a modest subset of what is possible using this approach. For instance, the CART algorithm used for model construction is amenable for extension to include both continuous and discrete multi-omics data if available without the need for feature normalisation. Embedded feature selection provided also ensures that the model only incorporates features that reduce uncertainty in the training set labels, such that a large feature space may be explored and reduced. In addition to incorporating other feature information, this tree-based approach is agnostic to the developmental context and could be applied to any step ontogenetic step where appropriately labelled samples are available. Lastly, we believe this approach holds significant promise for hypothesis and model generation as a natural result of the interpretable structure and information theoretic rule set learning.

Further, we have identified several avenues for future work that we believe show particular promise. Firstly, the eutelic nature of *C. elegans* provides an exciting opportunity to learn decision tree classifiers at each of every branching point during development, given appropriately sampled single-cell transcriptomic samples. As highlighted in section 2.3, this would allow for a complete mapping of development that builds on our knowledge of the lineage hierarchy to also learn and encompass the most informative transcriptomic features and thresholds. Secondly, we propose that decision tree classifiers could be trained on tumour sub-clones and the resulting models mined to identify key drivers of mutations. Thirdly, the information theoretic embedded feature selection could prove helpful for devising minimal GRNs of systems for which we have little prior understanding.

In conclusion, whilst these decision tree models appear simple we have shown how their learnt structure can begin to provide basic intuitions of complex developmental systems following the cursory model development highlighted here. Ultimately, these models performed well on withheld data, provide keen insights into the observed biological processes, present an exciting development of traditional computer science methods and fill a pressing gap in the toolbox of any statistically-minded biologist working with single-cell data to characterise developmental processes or for validating GRNs constructed through other means.

## 4 Methods

### 4.1 Model construction

#### 4.1.1 Recursive tree learning with CART

Finding the optimal binary decision tree amongst all possible topologies and combinations of constituent features and thresholds has been shown to be NP-complete (*30*). Without an efficient method for identifying the optimal tree, we could think of iterating through all possible trees. However, the complexity of this problem explodes with the tree depth and number of candidate features. In the increasingly common event where we have datasets of 100s or 1000s of genes, such an approach is intractable.

In light of this complexity, we turn to CART as a heuristic method that returns near-optimal models in computationally tractable time. This is achieved largely by substituting a globally optimal model for locally optimal splits at each successive branch in the tree. In such a top-down, greedy approach, trees are grown from a root node, and at each stage a feature is selected which is locally optimal with respect to some impurity measure - here Information Gain - at that point in the tree, regardless of whether such a branch might lead to a non-optimal model overall. Critically, whilst this approach is not guaranteed to return the globally optimum tree, the computation is cheap and the method shown to return near-optimal models in most cases (*31*).

The full set of labelled samples are divided into subsets according to a chosen feature. This feature is selected from the many others by evaluating how well it segregates labels in the resulting sub-populations. The process then recurses on the resulting daughter nodes until they each contain a single label or the nodes are deemed to be sufficiently pure, at which point the algorithm terminates and the final model is returned, Algorithm 4.1.1.

[H]

function growTree(node, D, depth) node.label = mode(*y*_*i*_: *i ∈D*) (*j*^*∗*^, *t*^*∗*^, *D*_*L*_, *D*_*R*_) = split(*D*) not worthSplitting(depth, cost, *D*_*L*_, *D*_*R*_) node node.left = growTree(node, *D*_*L*_, depth+1) node.right = growTree(node, *D*_*R*_, depth+1) node

The split function called in Algorithm 4.1.1 identifies the best feature, *j*^*∗*^, and best threshold for that feature, *t*^*∗*^, according to:

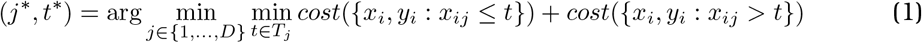

, where *T*_*j*_ is the set of possible thresholds for feature *j* obtained by sorting the unique values of *x*_*ij*_ (*17*) and the cost function corresponds to the chosen impurity measure - used to quantify the quality of a split with regards to purity of the daughter node labels (section 4.1.2).

For continuous features, such as the gene expression values derived by qPCR, ID3 considers thresholds at values for that feature already present in the training data (*32*). For each feature *j*, the cost function is calculated for all possible thresholds *t ∈T*_*j*_, corresponding to the unique expression values for gene j across all the cell samples in the training data. As shown in equation 1, the best attribute to split on is the gene and expression threshold (*j*^*∗*^, *t*^*∗*^) that maximises label purity over the proposed daughter nodes according to the chosen cost function. A visual summary of this tree building process is shown [Figure 2].

**Figure 1:**
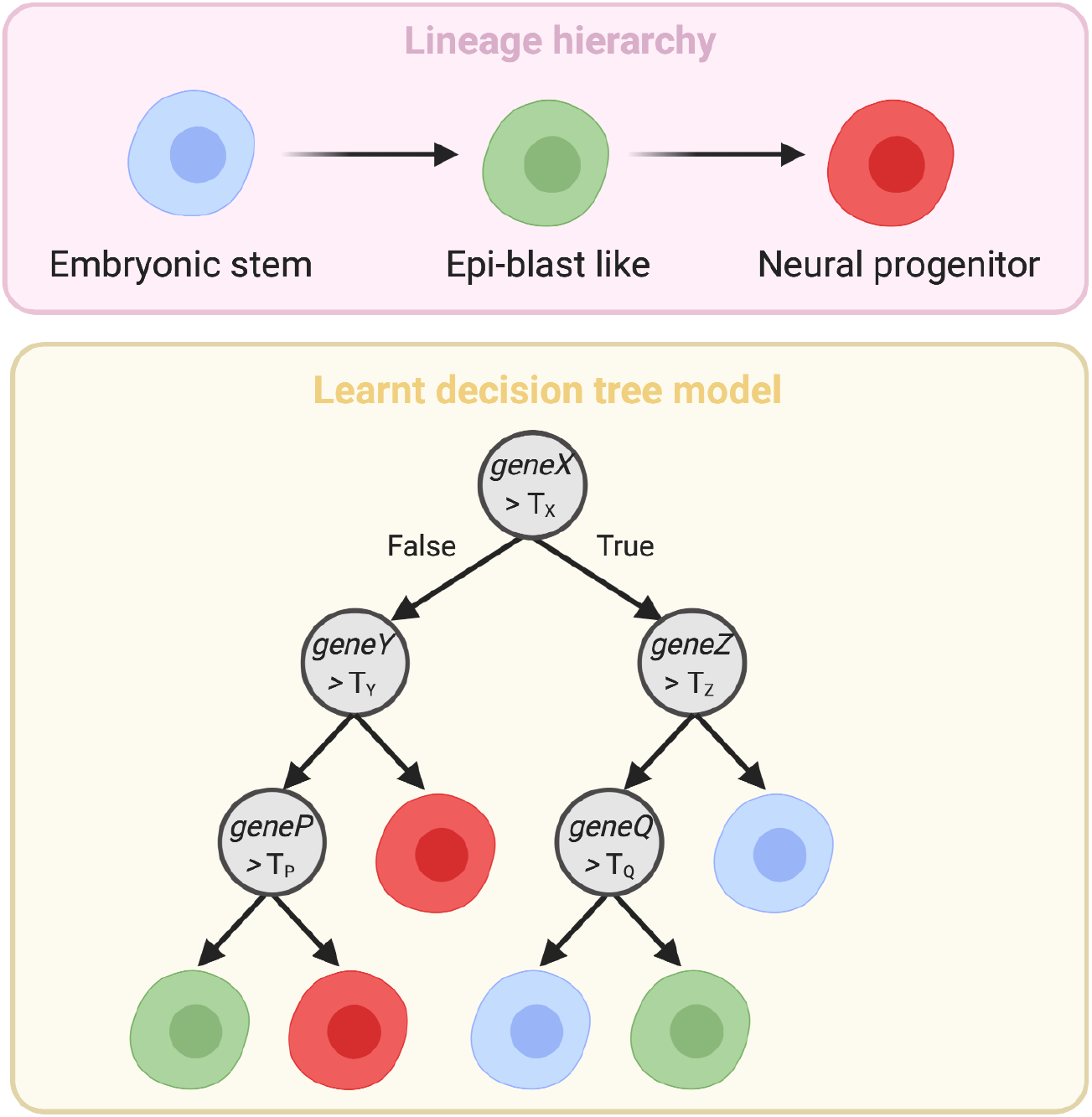
The simple lineage hierarchy for murine neuronal differentiation and a corresponding decision tree classifier. The learnt model structure showcases how the model assigns cell classes based on the information rich features (geneX, geneY, etc.) and corresponding expression thresholds (T_x_, T_y_, etc.) learnt during model training.

**Figure 2:**
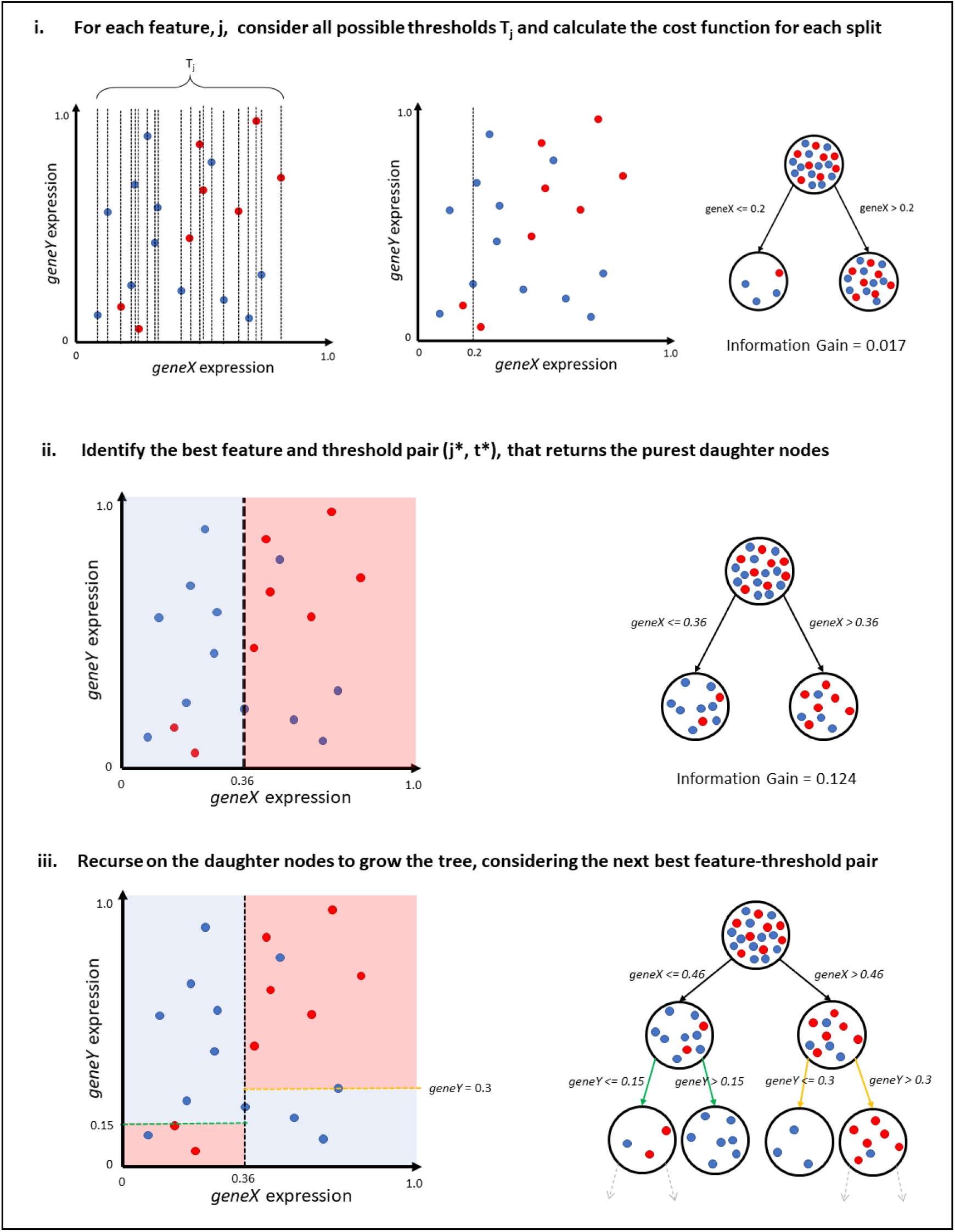
Decision tree construction protocol with the Classification and Regression tree (CART) algorithm for a binary classification task visualised in continuous two-dimensional feature space.

**Figure 3:**
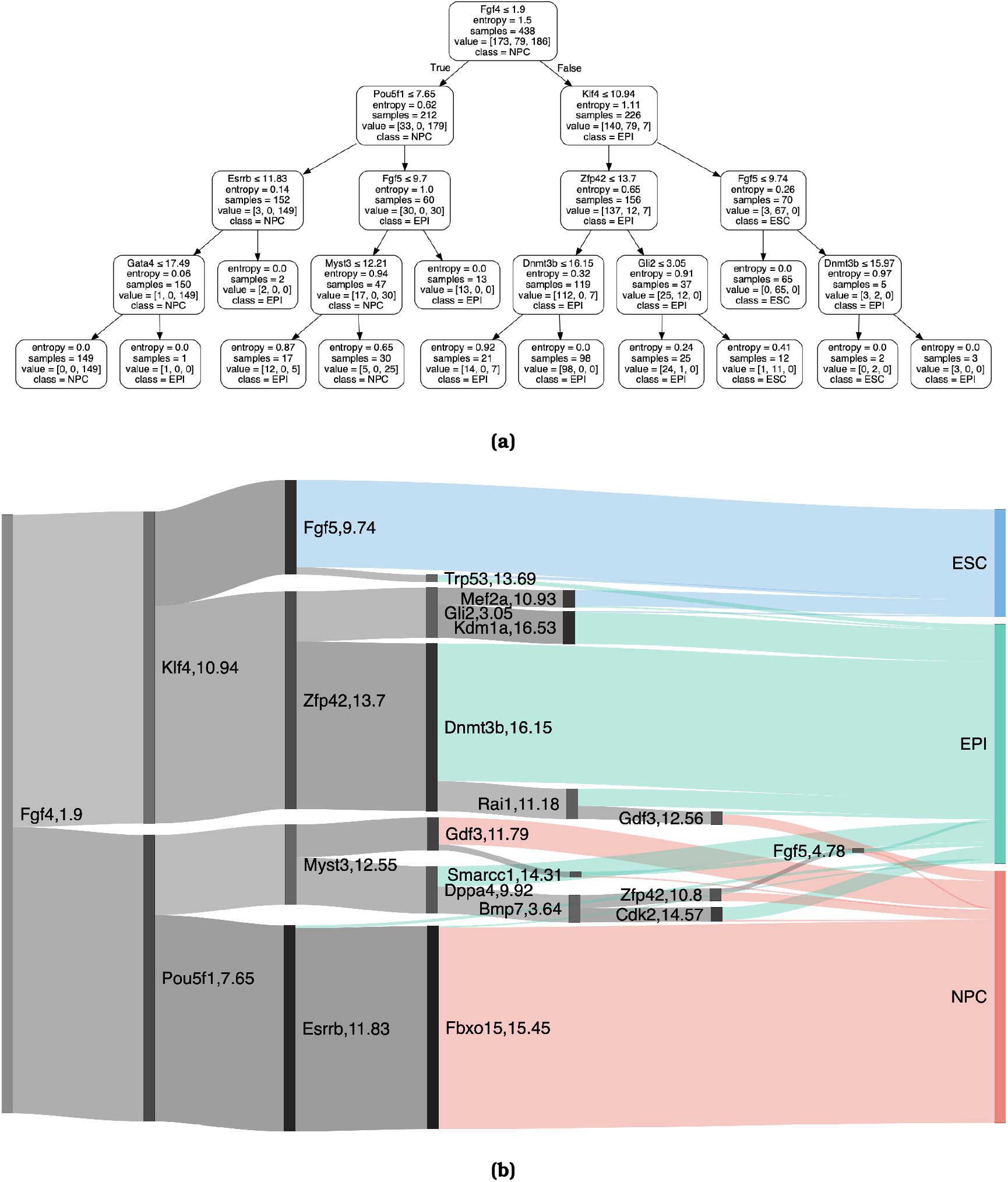
(a) Conventional representation of a decision tree model for the `Murine` data, trained on 80% of the full samples and limited in depth to four internal nodes. (b) Sankey representation of a decision tree classifier trained on a random 80% partition of the `Murine` data and shown to full depth. At each binary split, the gene feature is listed beside the learnt expression threshold.

**Figure 4:**
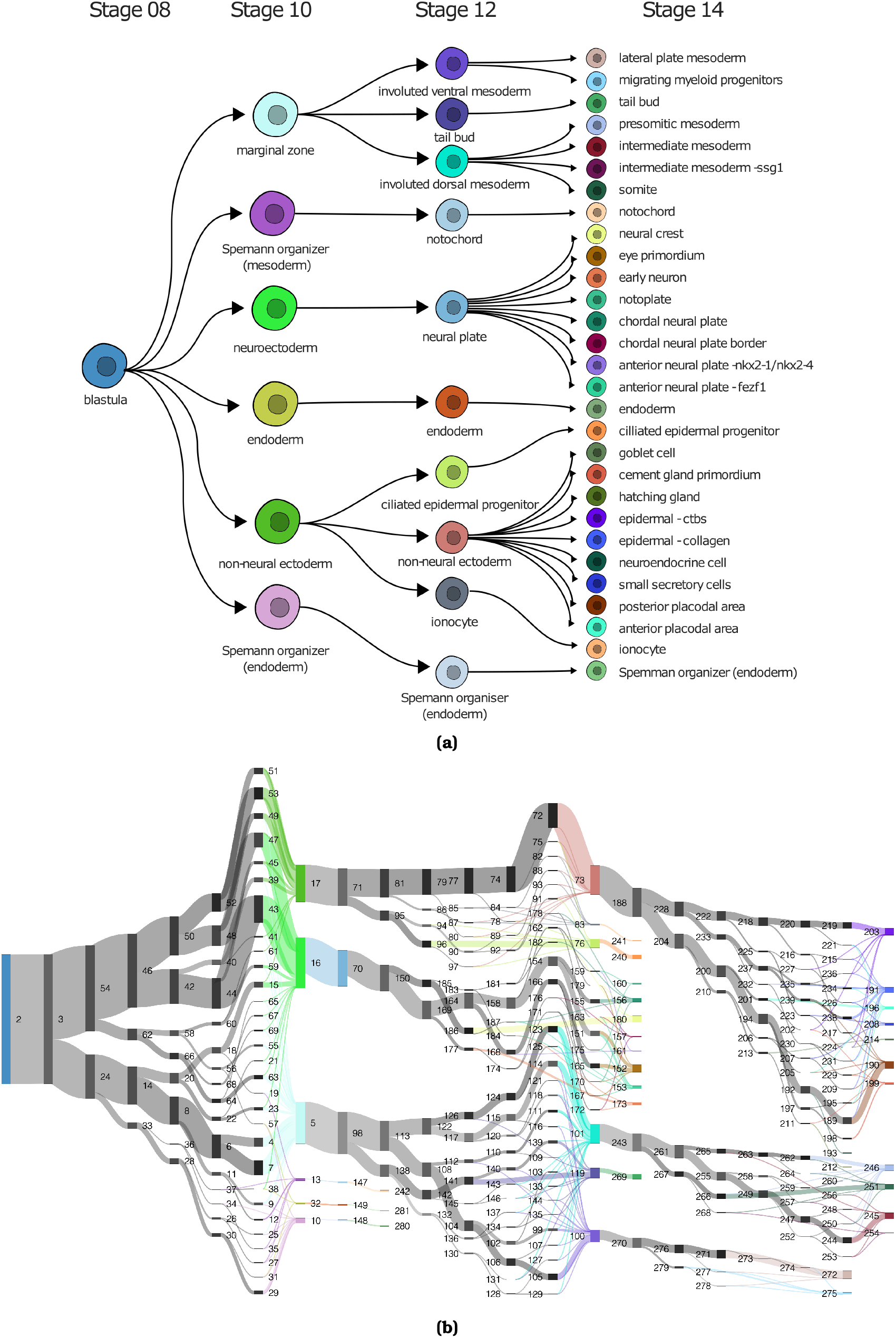
(a) Established lineage map for Xenopus tropicalis embryogenesis from stage 08 to stage 14 (7). (b) Decision tree models trained to distinguish the constituent cell classes at each branching point of the known lineage map. All nodes of the learnt models are numbered for legibility. The corresponding node information (features, thresholds, cell class) are provided in the supplementary material.

#### 4.1.2 Information Gain

Information Gain is a measure of the reduction in entropy in a data set given some transformation (*33, 34*). In our case, we calculate the information gain in the set of class labels, *S*, with respect to a candidate split, *A*, when a node is split into its left and right daughter nodes according to

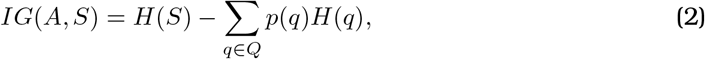

where *H*(*S*) is the entropy of the parent set S, Q is the set of subsets *q* proposed by splitting *S* on attribute *A* which thus satisfies *S* =⋃_*q∈Q*_ *q, p*(*q*) is the proportion of elements in the subset *q* compared to the set *S* and *H*(*q*) is the entropy over the subset *q*.

For clarity, the entropy of any set *S* containing *C* unique classes is

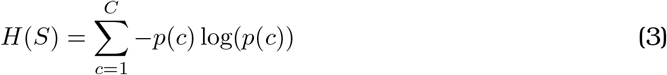

### 4.2 Model evaluation

#### 4.2.1 Misclassification rate and accuracy

After training our models, we may wish to evaluate the quality of their predictions on unseen data. Principally, we quantify the difference between the predicted labels and true (hidden) labels using the misclassification rate and accuracy. Let *ŷ*_*i*_ be the class prediction for the *i*^*th*^ sample in a test set of *n* samples. Then the test set misclassification rate is simply the fraction of misclassified cases, given by,

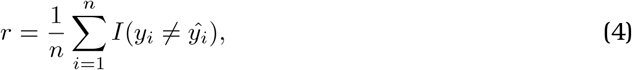

where *I* is the indicator function such that

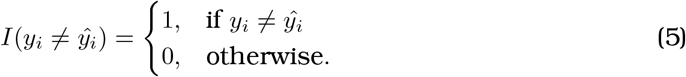

Following equation 4, the model’s accuracy on the test data is given by

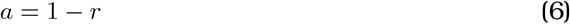

### 4.3 Minimal perturbations for sample reclassification

Consider a single cell whose expression profile for *J* genes is given by the vector **x** such that the gene expression value of the *j*th gene is given by **x**_*j*_: **x** = [*x*_1_, *x*_2_, …, *x*_*J−*1_, *x*_*J*_]. A learnt decision tree model, *M*, returns an estimate of the cell’s class label, *ŷ*. This classification, *M* (**x**) = *ŷ*, follows from the sample flowing through the tree from the topmost node to a leaf node whereby the sample is classified.

Given a new target class, *y*^*∗*^≠ *ŷ*, we seek to perturb **x** in such a way that the altered sample **x**^*′*^ is reclassified appropriately; *M* (**x**^*′*^) = *y*^*∗*^. We also hope to design this elementwise perturbation to the sample across all genes

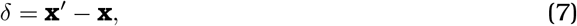

and we define the cost of applying such a perturbation as the summed absolute values over all genes as

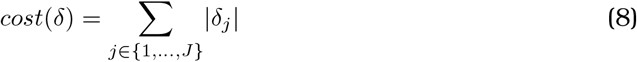

Each leaf node, *l*, in the set of leaf nodes, *L*, contained in model *M* has a unique path through the model, *P* (*l, M*), defining the sequence of nodes taken to reach *l* from the root node of *M*. For a leaf node at depth *z*, each internal node, *N*, contained in this path, *P* (*l, M*) = [*N*_1_, *N*_2_, …, *N*_*z−*1_, *l*] has a constituent feature id, *f* (*N*) *∈*{1, *J*}, as well as a corresponding expression threshold *t*(*N*) learnt during model training, and an associated direction, *d*(*N*) *∈ {*0, 1*}* given by

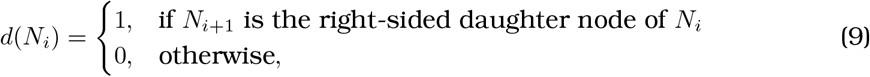

which describes whether the given path passes left or right at node *N* (returning 0 or 1, respectively).

Now, the perturbation required to set a sample onto the path of nodes defined by a given leaf node and keep it there can be represented by

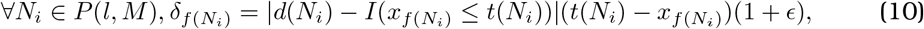

where *t*(*N*) is the threshold associated with each node *N* in the path, *x*_*f*(*N*)_ is the *f* (*N*)^*th*^ index of the sample *x* which contains the gene expression of gene *f* (*N*) for the sample *x, E* is a very small value increasing the magnitude of the distance between the sample’s value and threshold to successfully reroute the sample in cases where 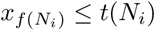, and *I*(*x*_*f*(*N*)_ *≤ t*(*N*)) is the indicator function such that

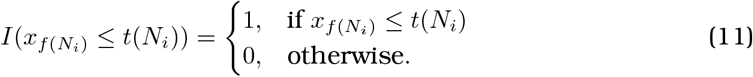

Note this ensures that the non-zero distance between the sample’s value for the feature and the associated threshold required for the sample to take the path defined by *P* (*l, M*) is only represented in *δ* if the unadulterated sample, *x* would otherwise stray from the path at that node, else 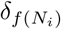 remains 0.

Thus, considering all leaf nodes with the desired target class, the perturbation with the smallest cost, *δ*^*∗*^, that if applied to sample *x*, given the model *M*, results in the appropriate reclassification is given by

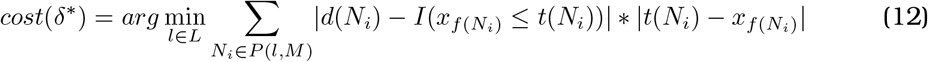

### 4.4 Datasets and preprocessing

#### 4.4.1 Murine data

The `Murine` data set contains single-cell gene expression data for mouse embryonic stem cells undergoing differentiation to neural progenitor cells. In the initial study, the mouse embryonic stem cells in the E14tg2a and R1 embryonic mouse cell lines were profiled during cell fate commitment, identifying a previously unknown epi-blast like, intermediary cell state (*20*).

In total, 547 cells were sampled over the course of seven days; at 24h, 48h, 72h, 120h and 168h after transfer from leukemia inhibitory factor (LIF) + 2i conditions to N2B27 neural basal medium; following existing protocol (*35*). Gene expression was reported for 96 genes affiliated with ES cell development using a high-throughput RT-PCR array. The resulting 547×96 feature matrix was supplemented by additional annotations provided in the initial work for each cell sampled. These included the sampling time for each cell, alongside a classification label for the cell; “ESC”, “EPI” or “NPC”; representing populations of embryonic stem cells, epiblast-like intermediate cells and neural progenitor cells respectively. As discussed in the supplementary material of the original work, these cell class labels were assigned through k-means clustering. Together with the gene expression data, these annotations provide us with a gold standard reference with which to train the models in our own work and to evaluate their predictive performance.

#### 4.4.2 Xenopus data

The `Xenopus` data is derived from a larger high-throughput sequencing effort, focused on early embryo development in *Xenopus tropicalis* (*7*). A total of 37,136 cells were sampled from zygotic genome activation (stage 8, 5 hours postfertilization) through to early organogenesis (stage 22, 22 hours postfertilization). Single cells were profiled with sc-RNAseq, following a previously established high-throughput single-cell droplet barcoding pipeline (*36*). The resulting data describes the underlying scRNA-seq counts for 25,087 genes in each of the sampled cells. Cell stage annotations were derived in the original work, using hierarchical clustering on the scRNA-seq data and cluster specific genes used to draw cluster labels from known embryonic cell types documented in the Xenopus Bioinformatics database - Xenbase (*37*). The full scRNA-seq count data and raw FASTQ files remain publicly accessible at https://kleintools.hms.harvard.edu.

For both the `Murine` and `Xenopus` data sets we have to acknowledge that there is a chance that the initial label assignment of cells may have been sub-optimal. Analyses choosing to incorporate decision tree models should take reasonable steps to ensure the accuracy of their labels in order for the downstream decision trees to be maximally informative.

### 4.5 Reproducibility

The complete code for training the models and deriving the results shown here has been made freely available at the cell-fate-decision-trees Github repository, https://doi.org/10.5281/zenodo.4342011

## A Supplementary material

### A.1 Rule set learnt by decision tree model

As discussed in section 2.2.2, here we provide the full Boolean rule-set learnt by the decision tree model to differentiate between ES, EPI and NP cells. As expected, there are 23 rules, corresponding to the 23 unique leaf nodes in the learnt model.

1. If Fgf4 ≤ 1.9 and if Pou5f1 ≤ 7.65 and if Esrrb ≤ 11.83 and if Fbxo15 ≤ 15.45 then sample belongs to class NPC
2. If Fgf4 ≤ 1.9 and if Pou5f1 ≤ 7.65 and if Esrrb ≤ 11.83 and if Fbxo15 > 15.45 then sample belongs to class EPI
3. If Fgf4 ≤ 1.9 and if Pou5f1 ≤ 7.65 and if Esrrb > 11.83 then sample belongs to class EPI
4. If Fgf4 ≤ 1.9 and if Pou5f1 > 7.65 and if Myst3 ≤ 12.55 and if Dppa4 ≤ 9.92 and if Bmp7 ≤ 3.64 and if Cdk2 ≤ 14.57 then sample belongs to class EPI
5. If Fgf4 ≤ 1.9 and if Pou5f1 > 7.65 and if Myst3 ≤ 12.55 and if Dppa4 ≤ 9.92 and if Bmp7 ≤ 3.64 and if Cdk2 > 14.57 then sample belongs to class NPC
6. If Fgf4 ≤ 1.9 and if Pou5f1 > 7.65 and if Myst3 ≤ 12.55 and if Dppa4 ≤ 9.92 and if Bmp7 > 3.64 and if Zfp42 ≤ 10.8 then sample belongs to class NPC
7. If Fgf4 ≤1.9 and if Pou5f1 > 7.65 and if Myst3 ≤12.55 and if Dppa4 ≤9.92 and if Bmp7 > 3.64 and if Zfp42 > 10.8 and if Fgf5 ≤ 4.78 then sample belongs to class NPC
8. If Fgf4 ≤1.9 and if Pou5f1 > 7.65 and if Myst3 ≤12.55 and if Dppa4 ≤9.92 and if Bmp7 > 3.64 and if Zfp42 > 10.8 and if Fgf5 > 4.78 then sample belongs to class EPI
9. If Fgf4 ≤1.9 and if Pou5f1 > 7.65 and if Myst3 ≤12.55 and if Dppa4 > 9.92 then sample belongs to class EPI
10. If Fgf4 ≤1.9 and if Pou5f1 > 7.65 and if Myst3 > 12.55 and if Gdf3 ≤11.79 then sample belongs to class NPC
11. If Fgf4 ≤ 1.9 and if Pou5f1 > 7.65 and if Myst3 > 12.55 and if Gdf3 > 11.79 and if Smarcc1 ≤ 14.31 then sample belongs to class NPC
12. If Fgf4 ≤ 1.9 and if Pou5f1 > 7.65 and if Myst3 > 12.55 and if Gdf3 > 11.79 and if Smarcc1 > 14.31 then sample belongs to class EPI
13. If Fgf4 > 1.9 and if Klf4 ≤ 10.94 and if Zfp42 ≤ 13.7 and if Dnmt3b ≤ 16.15 and if Rai1 ≤ 11.18 then sample belongs to class EPI
14. If Fgf4 > 1.9 and if Klf4 ≤ 10.94 and if Zfp42 ≤ 13.7 and if Dnmt3b ≤ 16.15 and if Rai1 > 11.18 and if Gdf3 ≤ 12.56 then sample belongs to class NPC
15. If Fgf4 > 1.9 and if Klf4 ≤ 10.94 and if Zfp42 ≤ 13.7 and if Dnmt3b ≤ 16.15 and if Rai1 > 11.18 and if Gdf3 > 12.56 then sample belongs to class EPI
16. If Fgf4 > 1.9 and if Klf4 ≤ 10.94 and if Zfp42 ≤ 13.7 and if Dnmt3b > 16.15 then sample belongs to class EPI
17. If Fgf4 > 1.9 and if Klf4 ≤ 10.94 and if Zfp42 > 13.7 and if Gli2 ≤ 3.05 and if Kdm1a ≤ 16.53 then sample belongs to class EPI
18. If Fgf4 > 1.9 and if Klf4 ≤ 10.94 and if Zfp42 > 13.7 and if Gli2 ≤ 3.05 and if Kdm1a > 16.53 then sample belongs to class ESC
19. If Fgf4 > 1.9 and if Klf4 ≤ 10.94 and if Zfp42 > 13.7 and if Gli2 > 3.05 and if Mef2a ≤ 10.93 then sample belongs to class ESC
20. If Fgf4 > 1.9 and if Klf4 ≤ 10.94 and if Zfp42 > 13.7 and if Gli2 > 3.05 and if Mef2a > 10.93 then sample belongs to class EPI
21. If Fgf4 > 1.9 and if Klf4 > 10.94 and if Fgf5 ≤ 9.74 then sample belongs to class ESC
22. If Fgf4 > 1.9 and if Klf4 > 10.94 and if Fgf5 > 9.74 and if Trp53 ≤ 13.69 then sample belongs to class ESC
23. If Fgf4 > 1.9 and if Klf4 > 10.94 and if Fgf5 > 9.74 and if Trp53 > 13.69 then sample belongs to class EPI

### A.2 Effect of tree depth and training-test partitions

Given heterogeneous training data, the structure of the learnt model and its resulting performance on unseen data can be affected by the initial random training/test partition. In addition, the maximum depth of a decision tree is a learnable hyperparameter with implications for how well the model can capture classification boundaries depending on the observed cell lineage. To demonstrate the influence of these factors on model performance, we generate 1000 random partitions of the data into 75% training and 25% test and for each partition we learn and evaluate decision tree models limited to various depths [Figure 6]. Whilst predictive performance is shown to vary across the 1000 models, the accuracy remains high for models grown to a depth of three internal nodes or more. As expected, models grown to a maximum depth of one internal node exhibit the poorest performance as they are only capable of allocating two of the three possible labels given the restricted topology. Again, trees learnt to a maximum depth of two internal nodes struggle to delineate the complex cell-fate decision boundaries. Increasing the maximum depth beyond 3 internal nodes for this simple lineage, we observe no significant improvements to model performance. Likely, any flexibility afforded to the learnt topology is countered by overfitting to the training samples.

**Figure 5:**
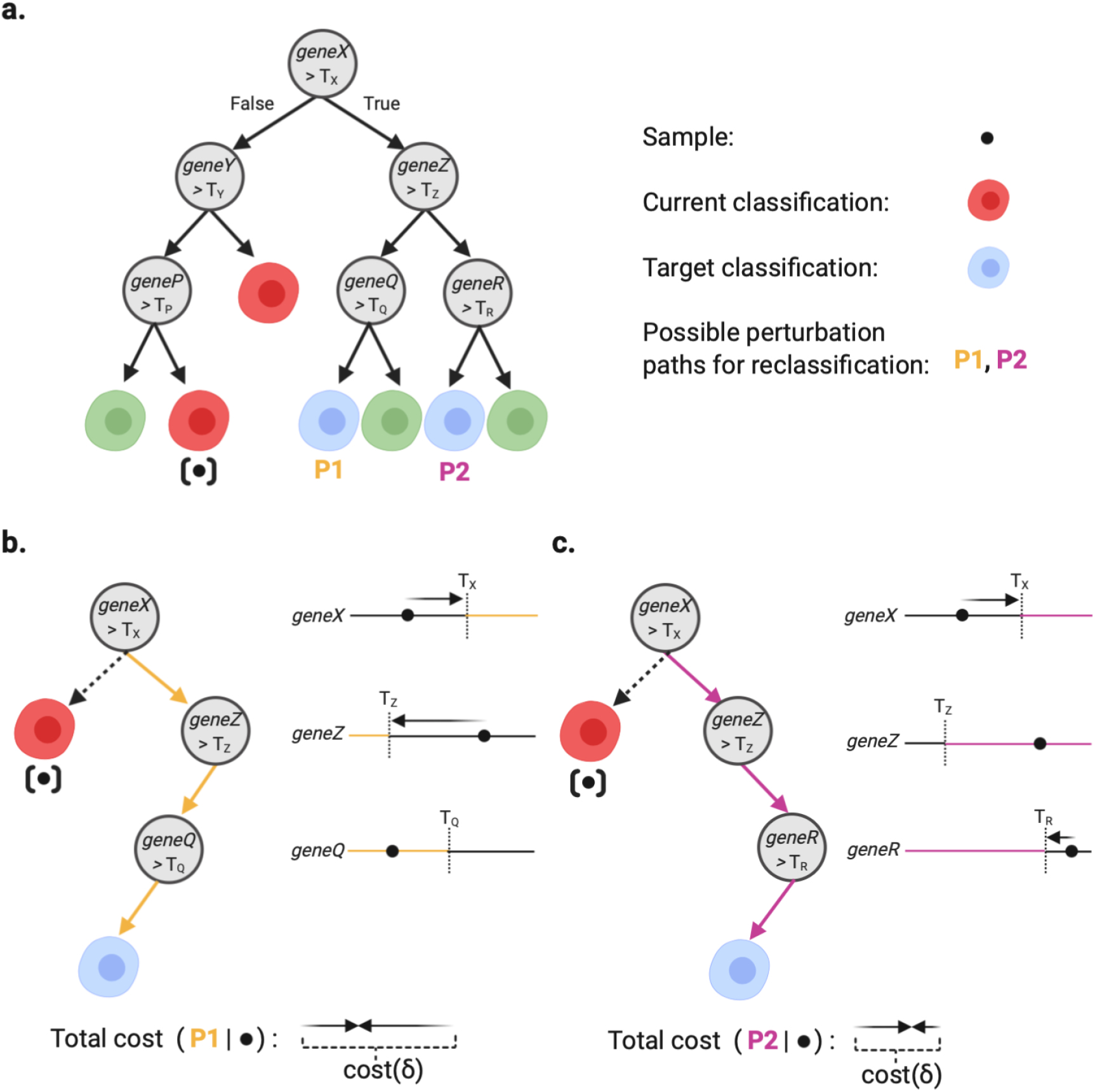
(a) Diagram showcasing the possible paths to targeted reclassification of a sample given a learnt decision tree model. The current classification of the sample is shown, as well as its terminal leaf node position in the learnt model. The two possible paths the sample could take in order for the sample to be appropriately reclassified by the model are signified by P1 and P2. (b) The path leading to the P1 leaf node is shown; drawn from the closest shared node of the P1 node and the sample’s original terminal leaf node (here, the root node). The minimal change to gene expression in order to force the sample down the P1 path is shown beside each node. Lastly the total cost of reclassifying the sample - in terms of summed absolute distance across the embedded features - is shown. (c) The equivalent schematic is also shown for the alternative path P2. We observe that for this example case the total cost of reclassification is lower for P2 than for P1, though taking either path will result in the sample being reclassified to the target class.

**Figure 6:**
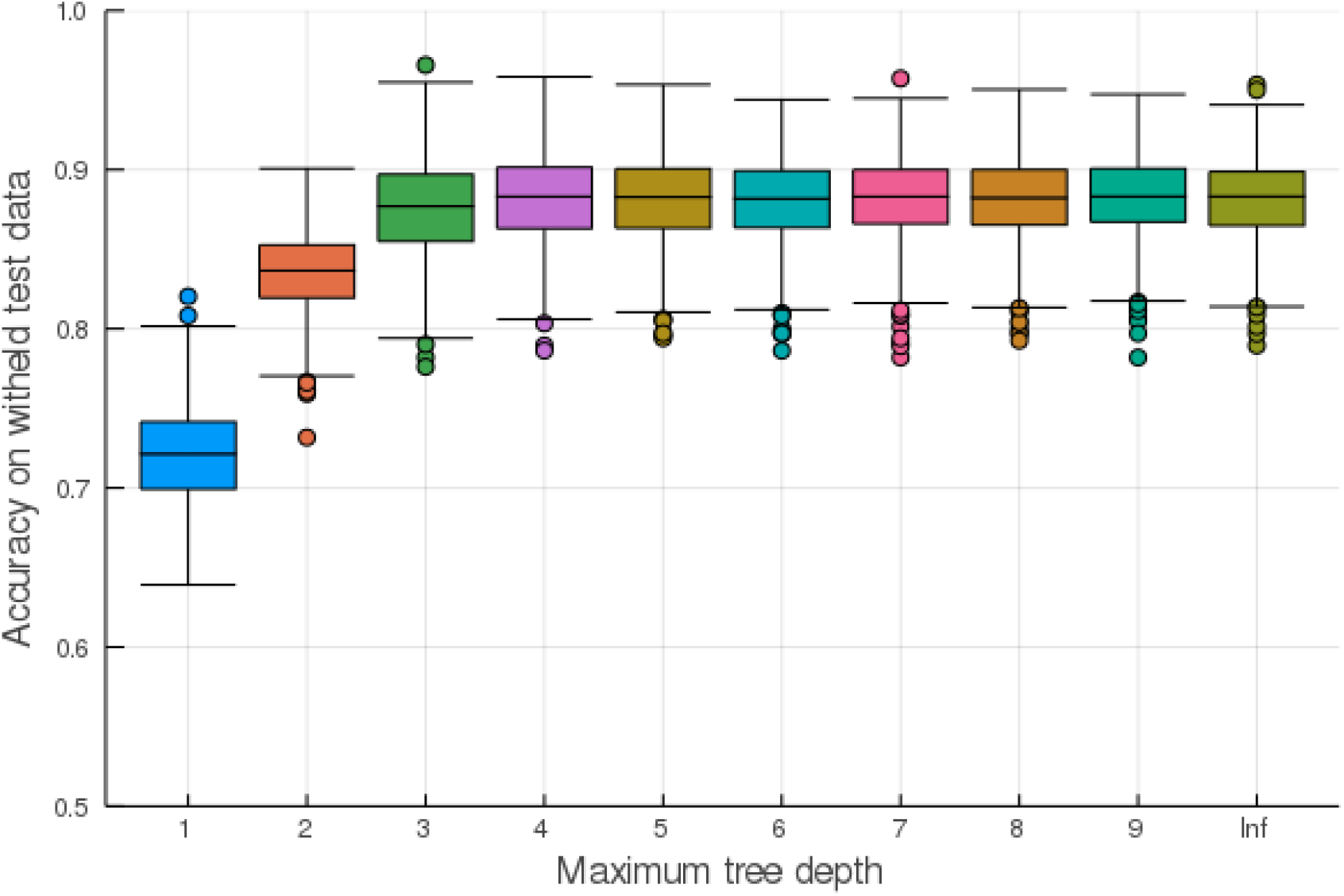
The performance of decision tree models is variant with the exact partition of data into training and test sets and is impacted by the maximum tree depth hyperparemeter. Here we show test set accuracy for models trained to specified maximum depths over 1000 random partitions of the training data into 75% training and 25% test sets.

